# SOCS-1 inhibition of type I interferon limits *Staphylococcus aureus* skin host defense

**DOI:** 10.1101/2020.09.28.317107

**Authors:** Nathan Klopfenstein, Stephanie Brandt, Sydney Castellanos, C. Henrique Serezani

## Abstract

The innate immune response to methicillin-resistant *Staphylococcus aureus* (MRSA) skin infection culminates in forming an abscess that prevents the bacterial spread and tissue damage. Pathogen recognition receptors (PRRs) dictate the balance between microbial control and tissue damage. Therefore, intracellular brakes are of fundamental importance to tune the appropriate host defense while preventing injury. The intracellular inhibitor suppressor of cytokine signaling 1 (SOCS-1); is a classic JAK/STAT inhibitor that prevents PRR responses by influencing the expression and actions of PRR adaptors and downstream effectors. Whether SOCS-1 is a molecular component of skin host defense remains to be determined. Here, we hypothesized that SOCS-1 decreases type I interferon production and IFNAR-mediated antimicrobial effector functions of the inflammatory response during MRSA skin infection. Our data show that MRSA skin infection enhances SOCS-1 expression, and both SOCS-1 inhibitor peptide treated and myeloid-specific SOCS-1 deficient mice display decreased lesion size, bacterial loads, and increased abscess thickness when compared to wild-type mice treated or not with scrambled peptide control. SOCS-1 deletion/inhibition increases phagocytosis and bacterial killing, dependent on nitric oxide release. SOCS-1 inhibition also increases antimicrobial effector function correlated with type I and type II interferon levels *in vivo*. IFNAR deletion and antibody blockage abolished the beneficial effects of SOCS-1 inhibition *in vivo*. Notably, we unveiled that hyperglycemia triggers aberrant SOCS-1 expression that correlates with decreased overall IFN signatures in the skin. SOCS-1 inhibition restores skin host defense in highly susceptible hyperglycemic mice. Overall, these data demonstrate a role for type I interferons in enhancing microbial clearance and host defense during MRSA skin infection.

## Introduction

*Staphylococcus aureus* is the leading cause of skin and soft tissue infections in the United States, accounting for almost 500,000 hospital admissions a year (Edelsberg et al., 2009). Although *S. aureus* colonizes ∼30% of the population, it is well suited to breaching the skin barrier, resulting in localized or potentially more severe systemic infections. The resistance of *S. aureus* to multiple antibiotics, particularly methicillin-resistant *Staphylococcus aureus* (MRSA), has made treatment of these infections increasingly difficult (Watkins et al., 2019). With the rise in antimicrobial resistance, there is a compelling need for safe, inexpensive, and non-antibiotic approaches that do not directly attack the bacterial target (which can lead to resistance over time), but instead host-centered strategies to prevent and treat these bacterial infections.

Skin resident macrophages, along with recruited monocytes and neutrophils, are responsible for major events during *S. aureus* skin infection, including recognizing the infection, abscess formation, and resolution of the inflammatory response (Cho et al., 2012; Cheng et al., 2011). Macrophages orchestrate abscess formation by establishing the inflammatory tone, recruiting neutrophils, killing microbes, clearing dead cells, and initiating wound healing. Since the abscess harbors viable and necrotic neutrophils plus bacteria at its core, it must be tightly organized to prevent deeper infection and bacterial dissemination (Brandt, Putnam, et al., 2018; Cheng et al., 2011).

Phagocytes contribute to host defense during multiple stages of *S. aureus* skin infection. While skin resident macrophages are involved in the initial recognition and killing of the bacteria, these cells are primarily engaged in producing chemoattractants to promote neutrophil and monocyte recruitment to the skin. Recruited neutrophils are crucial in *S. aureus* elimination through phagocytosis and killing via the generation of antimicrobial peptides, reactive oxygen (ROS) and nitrogen (RNS) species, as well as neutrophil extracellular traps (NETs) within the abscess (Brandt, Putnam, et al., 2018; Liu et al., 2018). Neutrophil-derived IL-1β also plays a critical role in optimal neutrophil recruitment to the site of infection, proper abscess formation, and improved infection outcome (Cho et al., 2012; Miller et al., 2006). Macrophages handle the resolution of the infection at the periphery of the abscess, which clears out the dead cell debris and breakdown the fibrous abscess capsule to allow for tissue healing and scar formation (Cheng et al., 2011; Santus et al., 2017).

Various pathogen recognition receptors (PRRs) recognize *S. aureus*, including scavenger receptors and the Toll-like receptors (TLRs) 1,2, 6, and 9 (Krishna & Miller, 2012). Both TLRs and IL1R utilize the adaptor protein myeloid differentiation primary response 88 (MyD88) for intracellular signaling (Feuerstein et al., 2015; Miller et al., 2006). MyD88 is a critical component of the host immune response to *S. aureus* skin infections (Brandt, Putnam, et al., 2018; Deguine & Barton, 2014). MyD88-deficient mice demonstrate impaired abscess formation and neutrophil recruitment during *S. aureus* skin, bone, and kidney infection, correlating with worse infection outcomes (Cho et al., 2012; Feuerstein et al., 2015; Miller et al., 2006). MyD88-dependent signaling culminates in activating different transcription factors such as NFκB, AP1, and IRFs (Deguine & Barton, 2014). Interestingly, TLR9 utilizes MyD88 to induce the production of type I interferons (IFNs). Furthermore, TLR9-mediated IFNβ production is essential to control *S. aureus* infection in the lungs.

Both type I and type II interferons are well-known enhancers of antimicrobial effector function in phagocytes. IFNγ increases host defense against a variety of pathogens (virus, bacteria, fungi, and parasites) (Dillen et al., 2018; Santus et al., 2017; Schroder et al., 2004), and the role of type I IFNs in the control of viral infections is well established while their role in bacterial infections has begun to emerge (Manca et al., 2005; Mancuso et al., 2007, 2009; Stanley et al., 2007). Recently, several manuscripts identified that type I IFNs can exert effects in the regulation of immune and tissue homeostasis upon bacterial insult that may have beneficial or detrimental consequences for the host (Carrero et al., 2004; Decker et al., 2005; McNab et al., 2015). *S. aureus*, as well as other pathogen-associated molecular patterns (PAMPs), enhance IFNα and IFNβ secretion *in vitro* and *in vivo* during infection via activation of intracellular PRRs (Mancuso et al., 2007). Interestingly, Kaplan et. al. have shown that *S. aureus* inhibits the expression of IFNβ in the skin and that treatment of mice with recombinant IFNβ decreases skin bacterial loads (Kaplan et al., 2012). Secreted IFNα and IFNβ binds to the heterodimeric interferon-alpha/beta receptor (IFNAR) and results in JAK1/Tyk2-mediated phosphorylation and dimerization of STAT-1, STAT-2, and IRFs (McNab et al., 2015).

STAT-1 activation is required for optimal skin host defense against *S. aureus*. However, STAT-1 enhances the expression of the JAK/STAT inhibitor suppressor of cytokine signaling-1 (SOCS-1) (Deguine & Barton, 2014; Takeda & Akira, 2004). We and others have shown that SOCS-1 inhibits STAT-1 dependent MyD88 expression in macrophages (Piñeros Alvarez et al., 2017; Serezani et al., 2011). We have also shown that SOCS-1 inhibits glycolysis and inhibits inflammatory responses during sepsis (Piñeros Alvarez et al., 2017). Whether SOCS-1 influences skin host defense remains to be elucidated. SOCS-1 acts through direct inhibition of the JAK tyrosine kinase to prevent STAT-1 phosphorylation and activation in a classical negative feedback loop (Liau et al., 2018). In addition to STAT-1, SOCS-1 also prevents the activation of different transcription factors, such as NF-κB and AP-1 (Naka & Fujimoto, 2010; Serezani et al., 2011; Yoshimura et al., 2007). SOCS-1 can also inhibit phagocyte function by 1) hampering the TLR-MyD88-dependent activation of NF-κB by targeting MyD88-adaptor-like protein (MAL) (Mansell et al., 2006); 2) inhibiting IL-1 receptor-associated kinase (IRAK) (Ahmed et al., 2015), 3) preventing MAPK signaling by binding to apoptosis signal-regulating kinase 1 (ASK-1) (Naka & Fujimoto, 2010). Since SOCS-1 exerts pleiotropic effects in phagocytes, it is expected that SOCS-1 could influence skin host defense. Indeed, SOCS-1 has demonstrated a detrimental impact in different infections, including viral, fungal, parasitic, and bacterial infections (Carrero et al., 2004; Decker et al., 2005; McNab et al., 2015). However, whether SOCS-1 is an important component of *S. aureus* skin host defense remains to be determined.

We hypothesized that SOCS-1 negatively impacts *S. aureus* skin infection outcomes by limiting the phagocyte dependent inflammatory response and host defense. Using a combination of *in vivo* bioluminescent imaging along with pharmacological (SOCS-1 blocking peptide) and genetic (myeloid-specific SOCS-1 deletion) approaches, we demonstrate that inhibition of SOCS-1 decreases lesion size and bacterial loads during *S. aureus* subcutaneous skin infection by increasing type I IFN-mediated macrophage antimicrobial effector functions.

Here, we identified a heretofore unknown role for SOCS-1 as a negative regulator of skin host defense in both homeostatic and hyperglycemic conditions. Our data also sheds light on the potential role of SOCS-1 inhibition as a host-directed therapeutic opportunity to treat antibiotic-resistant pathogens by redirecting the inflammatory response and increasing antimicrobial effector functions of phagocytes.

## Methods

### Animals

Mice were maintained according to National Institutes of Health guidelines for the use of experimental animals with the approval of the Indiana University (protocol #10500) and Vanderbilt University Medical Center (protocol #M1600215) Committees for the Use and Care of Animals. Experiments were performed following the United States Public Health Service Policy on Humane Care and Use of Laboratory Animals and the US Animal Welfare Act. C57BL/6J breeding pairs were initially obtained from the Jackson Laboratory and maintained by breeding at Vanderbilt University Medical Center (VUMC), Nashville, TN, USA. Wild-type BALB/c and IFNAR −/− mice were a gift from Dr. Stokes Peebles (VUMC). C57BL/6 *Socs1*^*fl*^ mice were obtained from Warren Alexander (Walter and Eliza Hall Institute, Parkville, Victoria, Australia) (Chong et al., 2003), and this strain was crossed with LysM^cre/cre^ mice (Jackson Laboratory) to create Socs1^fl^LysM^Cre^ (SOCS1^Δmyel^) mice. C57BL/6 *Socs1*^*fl*^ littermates were used as controls.

### Induction of hyperglycemia

For streptozotocin(STZ)-induced hyperglycemia, 6-to 8-week-old male C57BL/6J mice were treated by i.p. injection with 40 mg/kg of STZ (Adipogen) dissolved in 0.1 M sodium citrate buffer once daily for 5 consecutive days (Filgueiras et al., 2015). Euglycemic control mice received citrate buffer and served as vehicle control. Mice were considered hyperglycemic when blood glucose levels were >250 mg/dl. Mice were treated with STZ to induce hyperglycemia 30 days before MRSA skin infection.

### MRSA strains

The MRSA USA300 LAC strain was a gift from Bethany Moore (University of Michigan, Ann Arbor, Michigan, USA; ref. (Domingo-Gonzalez et al., 2013). The bioluminescent USA300 (NRS384 lux) strain was a gift from Roger Plaut (Food and Drug Administration, Silver Spring, Maryland, USA; ref. (Plaut et al., 2013)). The GFP-expressing USA300 strain was a gift from William Nauseef (University of Iowa, Iowa City, Iowa)(Greenlee-Wacker et al., 2014). MRSA stocks were stored at –80°C. MRSA was cultured as previously described (Dejani et al., 2016)

### MRSA skin infection and treatments

The murine skin infection model was adapted from a previous study (Dejani et al., 2016). Male mice between 6 and 12 weeks of age were used for MRSA skin infection. Mice were infected with approximately 3 × 10^6^ MRSA, and biopsies and sample collection were taken at various times, ranging from 6 hours to 9 days after infection, as previously described (Brandt, Klopfenstein et al., 2018; Dejani et al., 2016). Lesion size was measured daily via caliper, and the affected area was calculated using the standard equation for the area (length × width) (Becker et al., 2014). The iKIR (DTHFRTFRSHSDYRR) and scrambled-KIR peptide control (DTHFARTFARSHSDYRRI) were obtained from GenScript (Piñeros Alvarez et al., 2017). A lipophilic palmitoyl group was added to the N-terminus of both sequences to facilitate cell penetration (Waiboci et al., 2007). Mice were treated intraperitoneally with 50 μg of either the iKIR peptide or the scrambled peptide control (Piñeros Alvarez et al., 2017) 1 hour prior to infection and once-daily following infection. Mice were injected intraperitoneally with 40mg/kg of the anti-IFNGR or IFNAR antibody 3 hours before infection, followed by infection 24 hours.

### Skin Biopsy Specimens and Bacterial Load

8 mm punch biopsies were collected from naive and infected skin at different time points post-infection and used to determine bacterial counts, histological analysis, cytokine production, mRNA expression, and flow cytometry analysis (Brandt, Klopfenstein, et al., 2018). For bacterial counts, skin biopsy samples were collected, weighed, processed, and homogenized in tryptic soy broth (TSB) media. Serial dilutions were plated on tryptic soy agar. Colony-forming units (CFUs) were counted after incubation overnight at 37 °C and corrected for tissue weight. Results are presented as CFU/g tissue.

### Histopathology analysis

For histological analysis, 8 µm skin sections were stained with Hematoxylin and eosin or gram staining to visualize bacteria in the infected skin, as we have previously shown (Brandt, Klopfenstein, et al., 2018). Images of tissue sections were visualized and acquired using the Nikon Eclipse Ci and Nikon Ds-Qi2 (Nikon, Tokyo, Japan).

### In vivo bioluminescence imaging (BLI) and analysis with IVIS

An IVIS Spectrum/CT (Perkin Elmer) *in vivo* optical instrument was used to image bacterial bioluminescence in the mice. Bioluminescence imaging and analysis were performed as previously described (Brandt, Wang, et al., 2018).

### Skin Single Cell Isolation and Staining for Flow Cytometry

Skin biopsy specimens were collected and minced before digestion in 1mL of DMEM with 1 mg/ mL collagenase D (Roche Diagnostics) for 3 hours at 37°C. Single-cell suspensions were treated with CD16/32 Fc blocking antibodies (Biolegend; catalog 101310; clone 93) to prevent non-specific antibody binding and stained with the fluorescent-labeled antibodies for 20 minutes followed by fixation using 1% paraformaldehyde. The following antibodies were utilized: F4/80-FITC (Biolegend; catalog 123107; clone BM8), CXCR2-PE (R&D; catalog FAB2164P), Ly6G-PerCP/Cy5.5 (Biolegend; catalog 127616; clone 1A8) Ly6C-AF647 (Biolegend; catalog 128010; clone HK14), CD11b-PE/Cy7 (Biolegend; catalog 101216; clone M1/70). Analyses were completed using FlowJo software (FlowJo, Ashland, OR).

### Detection of Cytokines and Chemokines

Biopsy samples were collected, weighed, and homogenized in TNE cell lysis buffer containing phosphatase and protease inhibitors and centrifuged to remove cellular debris. Skin biopsy homogenates were then analyzed using the pro-inflammatory-focused 18-plex Discovery Assay from Eve Technologies (Eve Technologies, Calgary, AB) to detect cytokines and chemokines. Levels of IFNγIFNα, and IFNβ were measured the time points indicated in the legends by ELISA, (IFNγ -Invitrogen #88-7314) (IFNα-Biolegend #439407) (IFNβ-PBL #42120-1). Protein concentration was corrected for tissue weight.

### Immunoblotting

Western blots were performed as previously described (Serezani et al., 2011). Protein samples were resolved by SDS-PAGE, transferred to a nitrocellulose membrane, and probed with commercially available primary antibodies against SOCS-1, total-STAT-1, and phosphorylated STAT-1 (Y701) (all at 1:1000; Cell signaling), or β-actin (1:10,000; Invitrogen). Membranes were then washed and incubated with appropriate fluorophore-conjugated secondary antibodies (1:10,000, anti-rabbit IgG, IRDye 800CW antibody, #926-32211, Licor). Relative band intensities were quantified using ImageJ software (NIH), as previously described (Serezani et al., 2011).

### RNA Isolation and Quantitative Real-Time PCR

Skin biopsies samples were collected, and total RNA was isolated using lysis buffer (Buffer RLT; QIAGEN) following the manufacturer’s protocol. The RT^2^ First Strand Kit reverse transcription system (QIAGEN) was used for cDNA synthesis, and quantitative PCR (qPCR) was performed on a CFX96 Real-Time PCR Detection System (Bio-Rad Laboratories). Relative gene expression was calculated using the comparative threshold cycle (C_t_) and expressed relative to control or WT groups (ΔΔCt method). Primers for β-actin *and Socs1* were purchased from Integrated DNA Technologies (IDT, Coralville, IA).

### Nanostring and Gene Enrichment Analysis

Global gene expression in infected skin biopsies from wild-type hyperglycemic and euglycemic mice was assessed by the NanoString nCounter gene expression system (Nanostring, Seattle, WA). Mouse Myeloid Innate Immunity panel for 770 genes in 19 different pathways was used for the analysis. Designed CodeSet underwent extensive quality control to avoid cross-hybridization to non-target molecules in samples. RNA was extracted from the infected skin using Trizol. The purity and concentration of the RNA were confirmed spectrophotometrically with a Nanodrop (Thermo Scientific, Waltham, MA). 100 ng of RNA (20ng/µl) was hybridized with probe CodeSet before running samples on NanoString. nSolver 3.0 software used to assess the quality of the run, followed by the deletion of the low-quality sample from further analysis. Two-sided hypergeometric statistical analysis was performed with the Kappa Score threshold setting of 0.3. Enrichment depletion was calculated based on Benjamini-Hochberg Correlated p-values of <0.05.

### Bone Marrow-Derived Macrophage Generation

Bone marrow cells were flushed from both tibias and femurs of mice with ice-cold PBS. Cells were centrifuged, and red blood cells were lysed using ACK buffer. Cells were adjusted to 1×10^6^ cells/mL in DMEM with FBS (5%), M-CSF (20ng/mL), and GM-CSF (10ng/mL) and maintained at 37°C with 5% CO_2,_ and the cell culture media containing M-CSF and GM-CSF was changed every 3 days until day 7 when the cell was fully differentiated.

### Phagocytosis and Killing Assays

Bacterial phagocytosis and killing were performed as previously shown (Brandt, Klopfenstein, et al., 2018). Briefly, BMDMs (2 × 10^5^/well) were plated into two individual 96-well plates with opaque walls and clear bottoms. Cells were pretreated with 10 μM iKIR or scrambled (SCR) KIR peptide for 1 hour before the addition of GFP-MRSA at a multiplicity of infection of 50:1 (Brandt, Klopfenstein, et al., 2018). Infected cells were incubated 1 hour to allow phagocytosis, and both plates were washed with warm PBS, and GFP fluorescence was measured on the first plate. The second plate was then maintained in PBS with SCR KIR or PBS with iKIR peptide and was incubated for another 2 hours for killing assays. To determine the role of NO in microbial killing, cells were treated with 50 μM of the iNOS inhibitor 1400W dihydrochloride (Tocris).To measure the intensity of intracellular GFP fluorescence, extracellular fluorescence was quenched with 500 μg/mL trypan blue, and the GFP fluorescence was quantified using a fluorimeter plate reader. Trypan blue served as a blank. A reduction in GFP fluorescence in the killing plate relative to the phagocytosis plate indicated bacterial killing.

### Statistical Analysis

Results are shown as a mean +/− SEM and were analyzed using GraphPad Prism 8.0 software (GraphPad Software, San Diego, CA). For comparisons between two experimental groups, a Mann-Whitney test was used, and for comparisons among three or more experimental groups, one-way ANOVA followed by Bonferroni multiple comparison test was sued. P < 0.05 was considered significant.

## Results

### SOCS-1 impairs skin host defense

To explore the role of SOCS-1 in *S. aureus* skin infection, we initially examined *Socs1* mRNA expression in the MRSA-infected skin of C57BL6/J mice. We observed a gradual increase in skin *Socs1* mRNA expression at days 1 and 3 post-infection (Fig. 1A). Next, we investigated the consequences of SOCS-1 inhibition in skin host defense using pharmacologic and genetic approaches. During infection with bioluminescent MRSA, wild-type mice were treated 1 hour before infection with the iKIR or SCR KIR peptide (Jager et al., 2011; Piñeros Alvarez et al., 2017; Waiboci et al., 2007) followed by daily treatments post-infection. Mice treated with iKIR showed decreased bacterial loads when compared to infected mice treated with the scrambled control KIR (Fig. 1B and 1C), as evidenced by decreased bacterial bioluminescence, as well as decreased CFUs (Fig. 1D). Interestingly, iKIR treatment decreased bacterial burden in the skin of mice as early as six hours post-infection that persisted over the course of a nine-day infection (Fig. 1D). Lower bioluminescent signals also correlated with a more compact abscess in iKIR-treated mice (Fig. 1E).

**Figure 1.).**
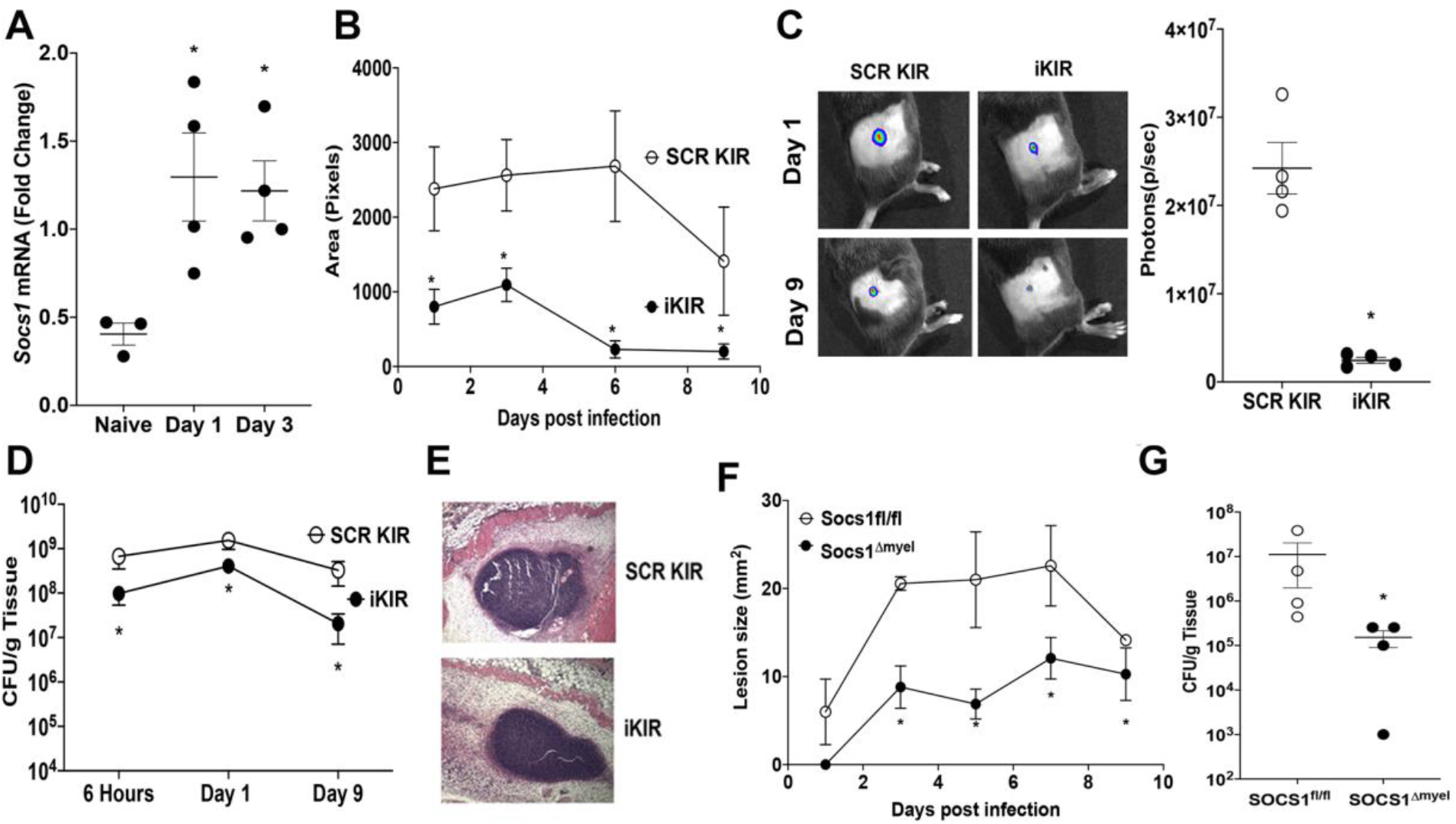
Inhibition of SOCS-1 actions improves subcutaneous skin infection outcome. **A)** mRNA expression of *Socs1* in the skin of mice infected subcutaneously with MRSA at day 1 and day 3 post-infection as determined by qPCR. **B)** Bioluminescent infection area in mice treated with SCR or SOCS-1 iKIR peptide using the *in vivo* animal imaging (IVIS Spectrum) detection system. **C) Right –** Total flux (photons/sec) of bioluminescent MRSA detected in mice treated as in **B** using IVIS Spectrum. **Left –** Representative images of bioluminescent MRSA in the skin of mice treated as in **B** using planar bioluminescent imaging. **D)** Bacterial load determined by CFU measured in the skin biopsy homogenates from mice treated as in **B** determined at the indicated time points after infection. **E)** Representative H&E stains from mice treated as in **B** and shown at 10X magnification **F)** Infection area measured every other day for 9 days post-infection in SOCS1^fl^ and SOCS1^Δmyel^ mice. **G)** Bacterial load determined by CFU measured in the skin biopsy homogenates in SOCS1^fl^ and SOCS1^Δmyel^ mice at day 9 post-infection. Data represent the mean ± SEM from 3–5 mice. *p < 0.05 vs. SCR KIR.

SOCS-1 regulates both the adaptive and innate immune response (Liau et al., 2018). As neutrophils and macrophages are required to control MRSA skin infection (Brandt, Putnam, et al., 2018; Liu et al., 2018), we further studied the role of myeloid-specific SOCS-1 actions during skin infection. Infection in *Socs1*^Δmyel^ mice resulted in smaller lesion size over time and decreased bacterial burden in the skin at day 9 post-infection (Fig. 1F and G). Together, these data suggest that phagocyte SOCS-1 expression negatively impacts MRSA skin infection outcomes.

### SOCS-1 inhibition increases bacterial phagocytosis and killing in macrophages

Due to the impact of SOCS-1 inhibition on bacterial clearance in the skin, we hypothesized that SOCS-1 might inhibit macrophage antimicrobial effector functions, such as phagocytosis and bacterial killing. We next examined Gram stains in the infected skin sections of wild-type mice treated with iKIR or scrambled KIR to determine whether differences in bacterial ingestion, niche location, and burden were evident between iKIR-treated or scrambled KIR-treated animals. MRSA was found mostly within cells in iKIR-treated mice, while a higher abundance of extracellular bacteria was observed in the skin of scrambled KIR-treated animals at day 1 after infection. (Fig 2A). When BMDMs from both *Socs1*^Δmyel^ mice and wild-type mice treated with the iKIR peptide were challenged with GFP-MRSA, we observed enhanced bacterial phagocytosis when compared to *Socs1*^*fl*^ BMDMs, or SCR KIR treated cells (Fig 2B). Importantly, SOCS-1 inhibition enhanced bacterial killing in BMDMs (Fig 2C). These data suggest that SOCS-1 is a negative regulator of macrophage antimicrobial effector functions that might be involved in MRSA skin infection control.

**Figure 2).**
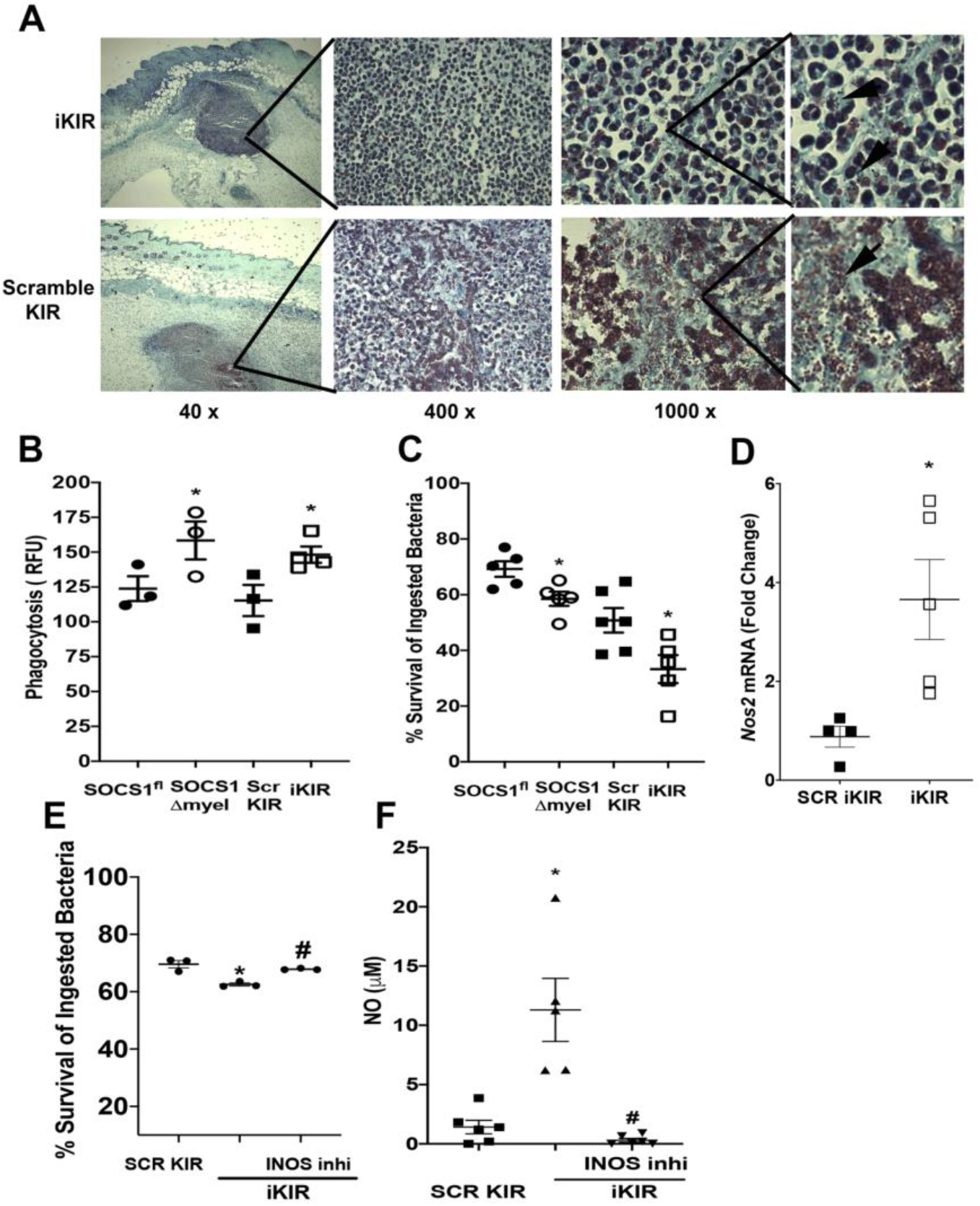
SOCS-1 inhibition enhances antimicrobial effector functions in BMDMs. **A)** Gram staining of skin biopsies collected at day 1 post-MRSA skin infection from iKIR and scrambled KIR treated mice. Gram staining to label gram-positive bacteria is shown in purple/brown. Magnifications are as shown. Black arrows indicate extracellular MRSA clusters. Images are representative of 3–5 mice per group. **B)** Phagocytosis of GFP tagged MRSA by BMDM’s from SOCS1^fl^ and SOCS1^Δmyel^ mice or BMDM’s from WT mice treated with the SCR KIR or iKIR peptide. **C)** Determination of Bacterial killing of GFP tagged MRSA by BMDMs from **B** as described in the Materials and Methods. **D)** mRNA expression of *Nos2* in the skin of infected mice treated with either SCR KIR or iKIR peptide at day 1 post-infection as determined via qPCR. **E)** Determination of bacterial killing of GFP-tagged MRSA as in **C** with BMDMs from WT mice treated with either SCR KIR, iKIR, or iKIR+ an iNOS inhibitor (1400W dihydrochloride). **F)** Measurement of nitric oxide in the supernatant of BMDMs from **D**. Data represent the mean ± SEM from 3–5 mice. *p < 0.05 vs. SOCS1^fl^ or SCR KIR treated mice. #p<.05 vs. iKIR treated mice.

*S. aureus* is sensitive to killing via nitric oxide (NO) (Brandt, Putnam, et al., 2018; Krishna & Miller, 2012; Zhang et al., 1994). To identify if SOCS-1 targets NO production in macrophages, we initially determined *Nos2* expression (the gene that encodes the inducible nitric oxide synthase protein) (iNOS) in iKIR treated and infected mice. We found significantly higher *Nos2* expression and NO production in the skin of iKIR-treated animals than mice treated with the scrambled peptide control at day 1 post-infection (Fig. 2D). As increased NO release could lead to more efficient bacterial killing *in vivo*, we determined whether increased iNOS/NO in iKIR treated mice accounts for improved microbial killing. BMDMs were treated with iKIR and an iNOS inhibitor (1400W dihydrochloride), followed by bacterial killing determination. iNOS inhibition ablated the beneficial effects of SOCS-1 inhibition on bacterial killing (Fig 2E). Notably, SOCS-1 inhibition increased NO release in MRSA-challenged BMDMs, and the treatment with the iNOS inhibitor reversed this trend (Fig 2F). Together these data demonstrate a role for SOCS-1 in regulating antimicrobial programs in macrophages dependent on NO release.

### SOCS-1 inhibits STAT-1 dependent production of pro-inflammatory cytokines and neutrophil recruitment during skin infection

Next, we examined the impact of SOCS-1 inhibition on the inflammatory milieu during skin infection. Initially, we determined whether SOCS-1 inhibition influenced STAT-1 activation *in vivo*. We found that while *S. aureus* infection induces STAT-1 phosphorylation (Y701), iKIR treatment further enhanced STAT-1 phosphorylation in the infected skin compared to mice treated with the scrambled KIR control (Fig 3A and 3B). Importantly, iKIR did not change SOCS-1 expression (Fig. 3A and 3B). We also confirmed that skin infection in *Socs1*^Δmyel^ mice increased pSTAT-1 abundance compared to infected WT control animals. These data suggest that the SOCS-1/STAT-1 signaling axis is a component of the skin host defense.

**Figure 3.).**
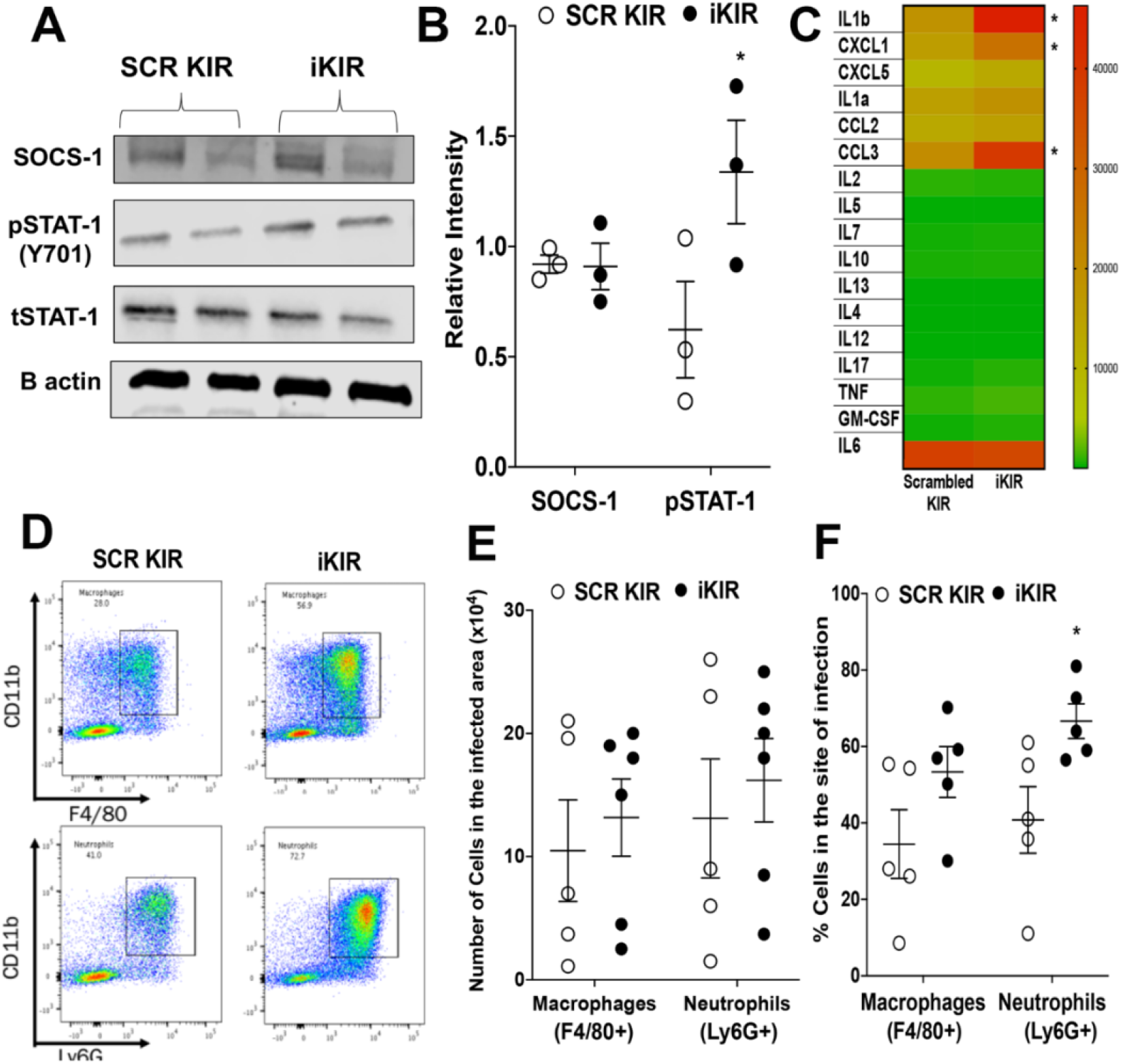
SOCS-1 inhibition increases pro-inflammatory cytokines and immune cell recruitment during skin infection. **A)** Representative Western blot for SOCS-1, PSTAT-1, and tSTAT-1 from biopsies collected at day 3 post-infection in SCR KIR and iKIR treated animals **B)** Densitometry quantification of 3 separate Western Blots as shown in **A. C)** Heat-map of proteins involved in the inflammatory immune response and its resolution in mice treated with either iKIR or SCR KIR at day 1 post-infection as measured using bead array multiplex (Eve Technologies). Proteins are listed on the left-hand *y*-axis, grouped alphabetically in clades. Red indicates higher abundance, whereas green represents lower abundance. Each column for each condition represents a technical replicate (*n* = 4-5/group). **D)** Representative flow plots from biopsies at day 1 post-infection in SCR KIR and iKIR treated mice for macrophages (F480+ CD11b+) and Neutrophils (Ly6g+ CD11b+). **E)** Total number of macrophages (CD11b+, F4/80+) and neutrophils (CD11b+, Ly6g+) in skin biopsies from **D. F)** Percentages of macrophages and neutrophils from biopsies in **D**. Data represent the mean ± SEM from 3–5 mice. *p < 0.05 vs. SCR KIR treated mice.

We then determined whether iKIR enhances the production of other STAT-1 dependent and – independent cytokines and chemokines during skin infection. We analyzed the abundance of 17 cytokines/chemokines known to modulate the inflammatory response and microbial clearance. Our data show that iKIR enhances specifically the production of pro-inflammatory cytokines known to drive proper abscess formation and improved infection outcome (IL-1β) (Cho et al., 2012), neutrophil recruitment (CXCL1) (Su & Richmond, 2015), and monocyte recruitment (CCL2) (Gillitzer & Goebeler, 2001) in response to MRSA skin infection (Fig 3C). These data suggest that SOCS-1 controls a specific group of cytokines known to drive a particular and targeted host immune response to MRSA skin infection.

Since IL-1β, CXCL1, and MIP-2 can all drive neutrophil and monocyte recruitment during skin infection (Gillitzer & Goebeler, 2001), we aimed to investigate whether iKIR treatment increases neutrophil and/or monocyte-derived macrophage migration to the site of infection. We performed flow cytometry on skin biopsies collected from iKIR-treated and infected mice using specific antibody markers for both macrophages (CD11b+ F4/80+) and neutrophils (CD11b+ Ly6g+) (Fig. 3D). While we did not see any differences in total neutrophil or macrophage numbers between the two treatment groups (Fig. 3E), we detected a significant increase in the percentage of neutrophils but not macrophages within the infected skin of iKIR-treated mice (Fig. 3G). These data suggest that SOCS-1 is an endogenous inhibitor of immune cell recruitment by hampering the secretion of inflammatory chemoattractants necessary for host skin defense against MRSA.

### SOCS-1/type I interferon axis mediates skin host defense

SOCS-1 functions as part of a negative feedback loop between type I and type II IFN production and STAT-1 activation (Liau et al., 2018). Whether SOCS-1 regulates the production of type I and type II IFNs during skin infection remains to be determined. We detected higher IFNγIFNα, and IFNβ levels in iKIR-treated and infected skin biopsies compared to scrambled KIR-treated mice at day 3 post-infection (Fig 4A). Due to the prominent role of IFNγ in enhancing phagocyte antimicrobial effector functions, we first blocked IFNγ actions in iKIR-treated and infected mice using an anti-interferon gamma receptor (IFNGR) antibody or IgG control antibody before infection and treatment with either iKIR or scrambled KIR peptide. Mice treated with the anti-IFNGR antibody plus scrambled KIR showed higher bacterial burden than mice treated with the control antibody as early as day 1 post-infection (Fig. 4B). However, while iKIR plus IgG control decreased bacterial load, the treatment of mice with anti-IFNGR antibody did not prevent iKIR effects on bacterial clearance, demonstrating that IFNγ is not solely involved in the inhibitory effects of SOCS-1 during MRSA skin infection (Fig 4B).

**Figure 4).**
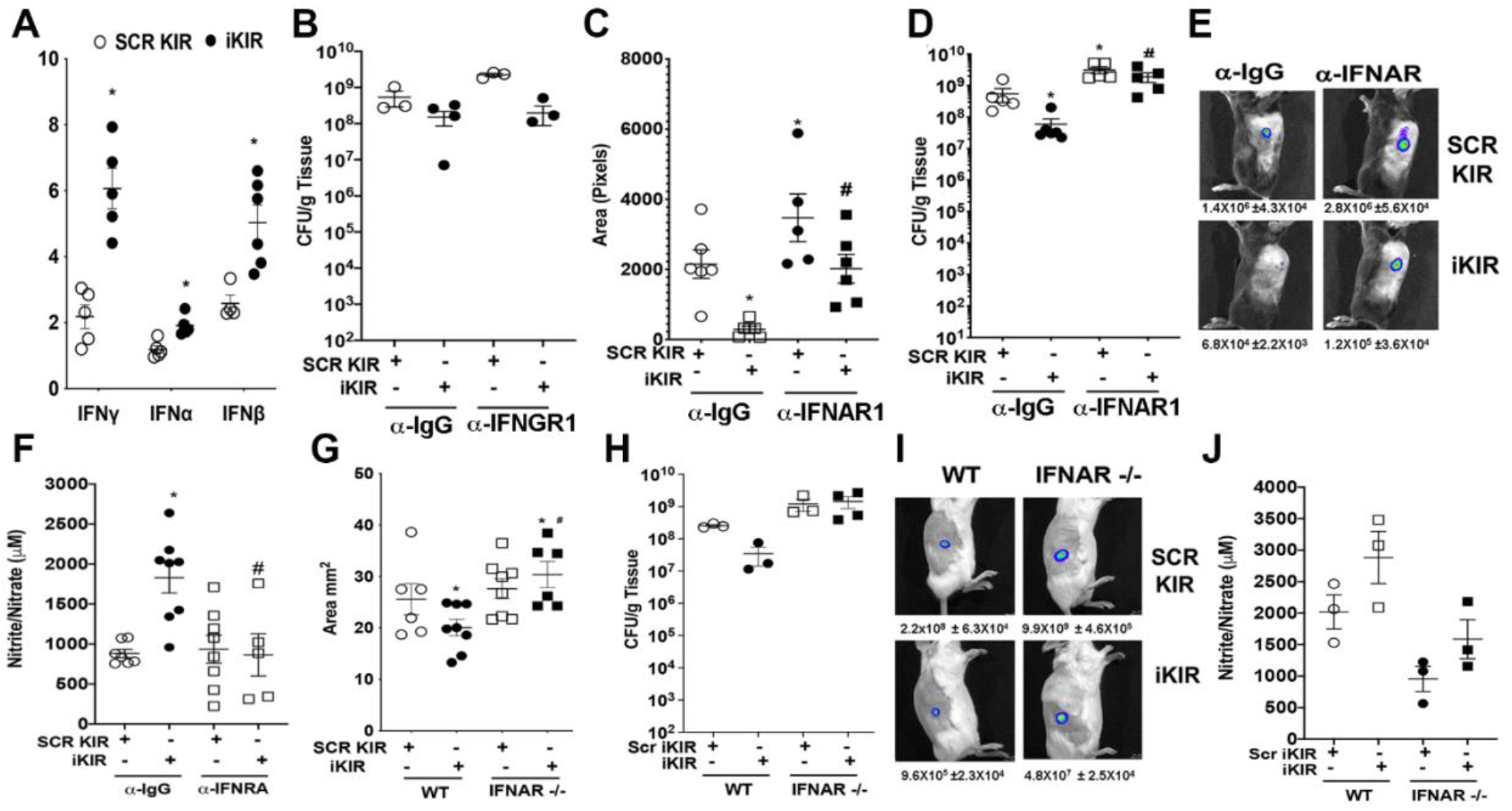
Inhibition of type I interferon signaling removes the benefit of SOCS-1 inhibition. **A)** Interferon levels in the skin at day 1 post-infection in SCR KIR and iKIR treated animals as measured via ELISA. **B)** Bacterial load as determined via CFU in skin biopsy homogenates of SCR KIR and iKIR treated mice treated with either IFNGR antibody or IgG control. **C)** Bioluminescent infection area in the skin of SCR KIR and iKIR treated animals treated with or without an IFNAR blocking antibody at day 1 post-infection. **D)** Bacterial load determined by CFU measured in the skin biopsy homogenates from mice treated as in **C** at day 1 post-infection. **E)** Representative images of bioluminescent MRSA in the skin of mice treated as in **C** using planar bioluminescent imaging with average flux (photons/sec) below **F)** Nitrite/Nitrate as measured via Griess assay from biopsies collected from mice treated as in **C** at day 1 post-infection. **G)** Surface lesion size as measured via caliper at day 3 post-infection in BALB/c or BALB/c IFNAR −/− mice treated with either iKIR of SCR KIR. **H)** Bacterial load determined by CFU measured in the skin biopsy homogenates from mice treated as in **G** at day 3 post-infection. **I)** Representative images of bioluminescent MRSA in the skin of mice treated as in **G** using planar bioluminescent imaging with average flux (photons/sec) below. **J)** Nitrite/Nitrate as measured via Griess assay from biopsies collected from mice treated as in **I** at day 3 post-infection. Data represent the mean ± SEM from 3–5 mice. *p < 0.05 vs. SCR KIR+αIGG or SCR treated WT mice. #p<0.05 vs. iKIR+ αIGG or iKIR treated WT mice.

Next, we determined whether iKIR actions are dependent on IFNα/β using both an IFNAR blocking antibody and IFNAR KO mice. While the pretreatment of mice with an anti-IFNAR antibody and scrambled KIR increased bacterial skin load, the blocking antibody also impaired iKIR-mediated decreases in lesion size (Fig 4C) and bacterial burden (Fig 4D and 4E). We then confirmed that type I interferons were a potential mediator of improved skin host defense during SOCS-1 inhibition in a genetic model. Wild-type and IFNAR^−/−^ mice were treated with either iKIR or scrambled KIR peptide, followed by MRSA skin infection. Our data showed that IFNAR^−/−^ mice had increased lesion size at day 3 post-infection (Fig 4G) and bacterial load compared to WT mice (Fig 4H and I). Furthermore, IFNAR^−/−^ mice were refractory to iKIR treatment (Fig. 4 G-I). Next, we aimed to link SOCS-1/type I IFN and NO in *in vivo* MRSA skin infection. Our data show that SOCS-1 inhibition enhances NO *in vivo* and blocking both SOCS-1 and IFNAR actions restored NO levels to the levels observed in scrambled KIR-treated mice (Fig. 4 F and J). Together, these data show a previously unknown operative axis of SOCS-1 and type I IFNs mediating NO-mediated microbial killing during bacterial skin infection.

### SOCS-1 drives impaired skin host defense in hyperglycemic mice

Patients with hyperglycemia are more susceptible to MRSA skin infections than euglycemic individuals (Chastain et al., 2019; Dejani et al., 2016). Therefore, we hypothesized that hyperglycemia enhances SOCS-1 expression and that blocking SOCS-1 actions might be a potential therapeutic approach to treat skin infections under hyperglycemic conditions. Initially, we determined *Socs1* mRNA expression over time in the skin of infected hyperglycemic and euglycemic mice. Our data show that *Socs1* expression was significantly upregulated in the skin of hyperglycemic animals at both day 1 and day 3 post-infection when compared to euglycemic animals (Fig. 5A). Enhanced SOCS-1 expression correlated with decreased pSTAT-1 in skin biopsies of infected hyperglycemic mice (Fig. 5B and 5C). Since SOCS-1 expression is increased in the skin of infected hyperglycemic mice, we asked whether the infection in hyperglycemic mice is characterized by an overall decrease in genes involved in IFN responses. Interestingly, our data show an overall downregulation of the IFN gene signature at day 1 post-infection in hyperglycemic mice (Fig. 5D). When we examined specific IFN-related genes, we observed inhibition of *Ifna1, Ifnb1, Ifng*, and *Stat1* in the infected skin of hyperglycemic mice. Importantly, we observed increased *Socs1* (confirming the results in Fig.4A and B), *Socs3, Irf2,Irf3*, and *Irf7* (Fig. 5E) in diabetic and infected mice. We then determined whether myeloid-specific SOCS-1 expression is involved in poor skin host defense in hyperglycemic mice. Our data show that hyperglycemic *Socs1*^*Δmyel*^ mice show decreased lesion size (Fig. 5F) and bacterial load (Fig. 5G) when compared to hyperglycemic *Socs1*^*fl*^ mice on day 9 post-infection. Next, we sought to investigate if iKIR could restore skin host defense and represent a potential therapeutic approach to treating antibiotic-resistant skin infections. Daily treatment of the hyperglycemic mice with iKIR significantly reduced lesions size and bacterial burden (Fig 5H and 5I). Together these data suggest that enhanced *Socs1* expression during hyperglycemia is detrimental to skin host defense and that iKIR treatment could represent host-directed immunotherapy to treat these antibiotic-resistant pathogens in the skin.

**Figure 5.).**
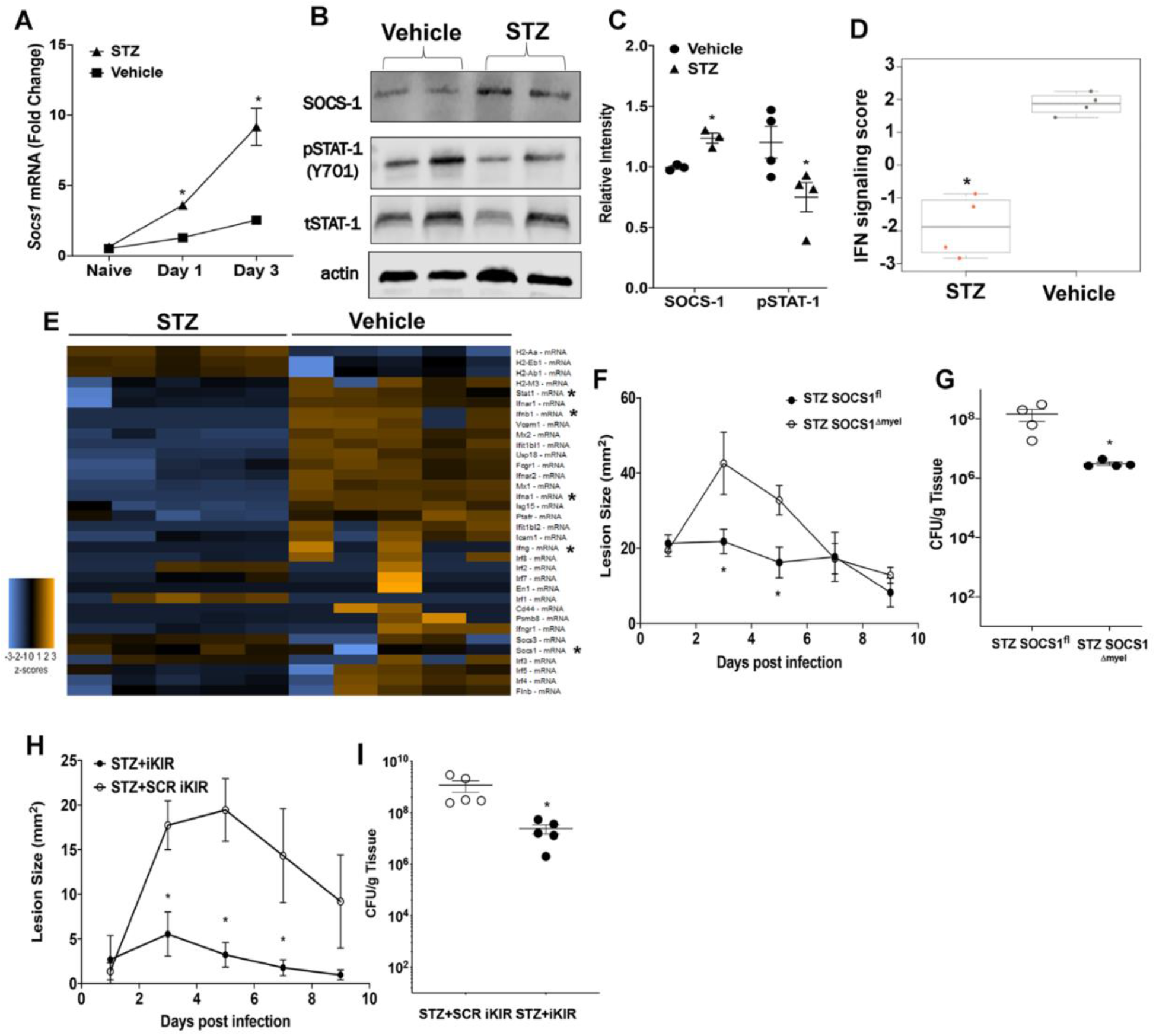
SOCS-1 impairs skin host defense in hyperglycemic mice. **A)** Expression of *Socs1* mRNA at day 1 and day 3 post-infection in biopsies collected from STZ induced hyperglycemic and control mice **B)** Representative Western blot for SOCS-1, PSTAT-1, and tSTAT-1 from biopsies collected at day 3 post-infection in STZ-induced hyperglycemic and control mice **C)** Densitometry quantification of separate Western Blots as shown in **B. D)** IFN signaling score determined using NanoSolver in mRNA isolated from infected skin of iKIR and scrambled KIR-treated mice. **E)** Heatmap of genes analyzed via NanoString from mRNA collected in **D. F)** Surface lesions size in STZ induced hyperglycemic and euglycemic SOCS1^fl^ and SOCS1^Δmyel^ mice over a 9 day subcutaneous MRSA infection. **G)** Bacterial load determined by CFU measured in the skin biopsy homogenates from mice treated as in **F** at day 9 post-infection. **H)** Surface lesions size in STZ induced hyperglycemic and euglycemic mice treated with either SCR KIR or iKIR over the course of a 9 day subcutaneous MRSA infection. **I)** Bacterial load determined by CFU measured in the skin biopsy homogenates from mice treated as in **H** at day 9 post-infection. Data represent the mean ± SEM from 3–5 mice. *p < 0.05 vs. Vehicle controls, SOCS1^fl^ or SCR KIR treated mice.

## Discussion

Monocytes, macrophages, and neutrophils are critical in the containment, elimination, and resolution of *S. aureus* infections in the skin and soft tissue. Due to the organ-specific diversity and pleiotropic function of these cells, by better understanding how these phagocytes respond to and eliminate pathogens, we might develop efficient therapeutic approaches to redirect the host’s immune response, improving microbial clearance while allowing for proper tissue repair. Host-directed immunotherapies are of particular importance, given the rapid rise of antibiotic resistance in several bacterial species, including *S. aureus* (Singer & Talan, 2014). Furthermore, the rise of community-acquired MRSA (CA-MRSA) in healthy individuals outside of the healthcare setting (Khan et al., 2018), further confirms the dire need for host-directed immunotherapeutic strategies for these infections.

Many studies have demonstrated the potential of host-directed therapies to treat skin infections. Few examples include the treatment of mice with antimicrobial peptides (Nizet et al., 2001), cytokines (IL-1β) (Miller et al., 2006), lipids (LTB_4_) (Brandt, Klopfenstein, et al., 2018), and enzyme inhibitors such as aspirin and indomethacin (Bystrom et al., 2008). Here, employing pharmacological and genetic approaches, we demonstrate that SOCS-1 inhibition/deletion boosts different arms of the skin immune response and increases MRSA clearance. SOCS-1 is a pleiotropic inhibitor of both the innate and adaptive immune response by controlling the actions of transcription factors and signaling effectors downstream of cytokine receptors, growth factors, and PRRs. We and others have shown that SOCS-1 inhibits MyD88 expression and actions (Serezani et al., 2011; Wu et al., 2015; Zahoor et al., 2020). SOCS-1 can prevent the differentiation of “pro-inflammatory” or M1 macrophages by limiting NF-κB p65 activation and STAT-1 actions (Duncan et al., 2017). Also, SOCS-1 inhibits TLR signaling by increasing MALD degradation and prevents MyD88 association with TLRs, thereby inhibiting the activation of downstream transcription factors (Liau et al., 2018). Therefore, given the role of TLRs and inflammatory cytokines in *in vivo* host defense, it is anticipated that endogenous SOCS-1 might help prevent tissue damage and, consequently, impacts phagocyte antimicrobial effector function. Indeed, infections with both gram-positive (Mancuso et al., 2007; Wu et al., 2015), gram-negative (Decker et al., 2005) bacteria as well as *Mycobacterium tuberculosis* (Manca et al., 2005) and parasites, such as *Leishmania major* (Bullen et al., 2003) and the fungus *Candida albicans* (Shi et al., 2018) promote SOCS-1 expression to actively suppress the immune response, allowing pathogen replication and immune evasion. Expression of SOCS-1 during infection correlates with reduced levels of the pro-inflammatory cytokines IL-1β, TNF-α, and IL-6 as well as antimicrobial NO and reactive oxygen species (ROS) (Duncan et al., 2017; Yoshimura et al., 2007). However, the mechanisms of SOCS-1 actions in infected phagocytes remains to be fully determined. Here, we advanced the field forward by demonstrating that myeloid-specific SOCS-1 inhibits phagocyte antimicrobial effector function, neutrophil migration to the site of the infection, and ultimately linked SOCS-1 actions to IFNα/β *in vivo*.

SOCS-1 is known as a STAT inhibitor, and therefore, most of the reported SOCS-1 actions might require transcriptional regulation and therefore take hours/days to be detected. Here, we unveiled a new role for SOCS-1 in the early macrophage response, namely phagocytosis and microbial killing. It has been shown that SOCS-1 deletion amplifies *C. albicans* phagocytosis by controlling IFNγ actions (Shi et al., 2018). Here, we separated the delayed transcriptional effects of SOCS-1 vs. early signaling effects using the iKIR peptide. Although we did not further investigate the intracellular targets of SOCS-1 in *S. aureus* phagocytosis, we identified NO as an intermediate of SOCS-1 actions in the microbial killing. A specific role for SOCS-1 in inhibiting phagocyte antimicrobial effector function was also evidenced by the detection of intracellular bacteria in the iKIR-treated mice, while in scrambled KIR treated mice, most of the bacteria were observed extracellularly. Our future studies will address this critical question, and we expect to unveil the targets and mechanisms involved in SOCS-1 inhibition of phagocytosis.

Chemokines and IFNs enhance both the recruitment and antimicrobial effector functions of phagocytes during infection (Gillitzer & Goebeler, 2001; Müller et al., 2018). Recent work has also demonstrated a novel role for IFNγ in the activation of the fibrinolytic system to control abscess thickness, improve leukocyte recruitment, and drive abscess breakdown during *S. aureus* and *C. albicans* infection (Santus et al., 2017). While IFNγ is a potent macrophage activator known to increase gene expression of many antimicrobial effectors such as members of the NADPH oxidase complex (gp91phox, p47phox), iNOS, and antimicrobial peptides, IFNγ also enhances the expression of MHCII and antigen presentation (Schroder et al., 2004). In our study, IFNγ production alone does not seem to account for the therapeutic benefit of SOCS-1 inhibition in skin host defense. Interestingly, SOCS-1 inhibition mimicked several aspects of the protective effects of IFNγ such as a highly organized and compact abscess with a thick capsule. If SOCS-1 regulates the expression of collagen and fibrinolytic proteins involved in abscess capsule formation remains to be determined.

Though the bulk of studies on IFNα and IFNβ focus on their role in anti-viral adaptive immunity, emerging studies demonstrate a role for IFNβ in macrophage activation and bacterial killing (McNab et al., 2015). IFNβ signaling in inflammatory macrophages has been shown to act in an autocrine signaling manner to promote iNOS expression and NO release along with other pro-inflammatory mediators to drive increased bacterial killing (Gao et al., 1998; Müller et al., 2018; Zhang et al., 1994). These studies highlight that while IFNβ treatment alone might not be enough to drive these phenotypes, IFNβ treatment combined with TLR1/2 and 4 stimulation significantly enhances NO levels. This fits into our current study since TLR2 is an important pathogen recognition receptor in binding the lipoteichoic acids that make up the MRSA’sl wall of MRSA (Müller et al., 2018). The autocrine nature of IFNβ signaling in macrophages would also explain why inhibition of SOCS-1 significantly improved infection outcomes at time points as early as 6 hours post-infection. Therefore, it seems likely that increased iKIR-dependent IFNβ production and/or actions during MRSA skin infection are promoting increased NO and bacterial killing in our current study. Interestingly, Kaplan et al. demonstrated that *S. aureus* infection in the skin inhibits IFNβ production and that treatment of mice with recombinant IFNβ increases bacterial clearance. Although we observed that MRSA enhances IFN α and β production in the skin, the significant differences between our manuscripts are the higher inoculum used in their work (1×10^7^) vs. 3×10^6^ used here, and the strain of S. aureus (Newman vs. MRSA USA 300). That different S. aureus strains exhibit different capabilities to induce IFN α/β production has been previously shown (Parker & Prince, 2012). If SOCS-1-mediated type I IFN actions are detrimental for other strains of *S. aureus* in different organs remains to be determined. Furthermore, a few more questions remain to be addressed. We did not specify whether SOCS-1 modulates IFNAR signaling directly, and acts on downstream events remain elusive. We speculate that SOCS-1 could be inhibiting the activation of different IRFs, such as IRF7 and IRF3, but more studies are needed to address further the potential mechanisms of SOCS-1 mediated IFNα/β production and/or signaling during infections. Our study shows that increased iKIR-mediated IFNβ production and/or IFNAR signaling is enough to promote sufficient bacterial killing in the absence of IFNGR signaling *in vivo*. However, due to the known role of IFNγ in macrophage activation, we cannot exclude a potential role for IFNγ in iKIR-treated mice, as we only determined microbial clearance in a single time point after infection.

To explore SOCS-1 inhibition as a potential therapeutic strategy in a highly susceptible population, we sought to investigate iKIR peptide improves host defense in hyperglycemic mice. We found that SOCS-1 expression is increased in macrophages from hyperglycemic animals. The iKIR peptide significantly decreased bacterial burden and lesion size compared to hyperglycemic mice treated with the scrambled peptide control. We also confirmed the beneficial role of SOCS-1 deletion in myeloid cells in skin host defense of hyperglycemic mice. Interestingly, we also unveiled an overall decrease in the expression of genes involved in IFN production and signaling. The basis for such drastic event is unknown, but it could include potential epigenetic modifications in the genes involved in IFN response. Nonetheless, these findings are intriguing and exciting, since we would expect that increased SOCS-1 expression in macrophages should decrease an overall exacerbated inflammatory response, but instead, we and others have shown that infection in hyperglycemic mice is characterized by a robust and detrimental migration of neutrophils and a delayed clearance of microbes (Brandt, Wang, et al., 2018; Dejani et al., 2016). In this context, aberrant intracellular SOCS-1 might be involved in preventing bacterial ingestion and killing while failing to regulate cytokine-dependent tissue inflammation. The mechanisms underlying SOCS-1 effects during infection in hyperglycemia/diabetes models are the current focus of our laboratory. Together this data suggests that SOCS-1 may be a potential target for future therapeutic intervention in skin infections and may benefit not only to highly susceptible patient populations but also to more broad patient populations with other bacterial infections of the skin.

## Author contributions

NK, SB, and HS conceived and designed the study; NK, SB, and SC conducted experiments. All authors contributed to data analysis. NK and HS wrote the manuscript, and all authors contributed to the manuscript editing.

## Conflict of Interests

The authors have declared that no conflict of interest exists.

### Acknowledgments

We thank Bethany Moore (University of Michigan, Ann Arbor, MI, USA) for providing the MRSA USA300 LAC strain and Roger Plaut (Food and Drug Administration, Silver Spring, MD, USA) for providing the bioluminescent USA300 (NRS384 lux) MRSA strain. We would like to thank the Serezani laboratory for the input. This work was supported by NIH grants R01HL124159-01 and RAI149207A (to CHS) and T32AI060519 (to SLB), and T32AI138932 (to NK).

